# Quantitative genetic study suggests sex-specific genetic architecture for fetal testosterone in a wild mammal

**DOI:** 10.1101/2022.02.18.480975

**Authors:** Ruth Fishman, Simona Kralj-Fišer, Sivan Marglit, Lee Koren, Yoni Vortman

## Abstract

Testosterone plays a critical role in mediating fitness-related traits in many species. Although it is highly responsive to environmental and social conditions, evidence from several species show a heritable component to its individual variation. Despite the known effects that in utero testosterone exposure have on adult fitness, the heritable component of individual testosterone variation in fetuses is mostly unexplored. Furthermore, testosterone has sex-differential effects on fetal development, i.e., a specific level may be beneficial for male fetuses but detrimental for females. The above mentioned may lead to a different genetic structure underlying the heritability of testosterone between the sexes. Here, we used a wild animal model, the feral nutria, quantified testosterone using hair-testing and estimated its heritability between parent and offspring from the same and opposite sex. We found that in utero accumulated hair testosterone levels were heritable between parents and offspring of the same sex. However, there was a low additive genetic covariance between the sexes, and a relatively low cross-sex genetic correlation, suggesting a potential for sex-dependent trait evolution, expressed early on, in utero.

## Introduction

Testosterone has a critical function in mediating fitness-related traits in both males and females (Hau 2007; Ketterson, Nolan, and Sandell 2005; Mills et al. 2009). Since it is involved in the development of male secondary sex characteristics and traits expressing male fertility status, high testosterone levels may be favored by sexual selection (Hillgarth et al., 1997). Selection can act on a trait if it varies among individuals, has a significant genetic base, and affects fitness (Falconer and Mackay, 1996). To understand the evolutionary implications of testosterone levels variation its heritability needs to be assessed (Boake, 2002). Testosterone shows high individual variance, and although it varies across life-history stages and environmental and social situations (e.g., Wingfield et al., 1990), it also shows a high degree of intra-individual repeatability (Kraus et al., 2020; Liening et al., 2010; Meikle et al., 1988; Mutwill et al., 2021; While et al., 2010; but see also Kralj-Fišer et al., 2007; Pavitt et al., 2015).

Male testosterone levels are positively associated with their reproductive success in several species (e.g., Adkins-Regan, 2005; Folstad and Karter, 1992; Mills et al., 2009; Negro et al., 2010). Testosterone levels have been linked to functional reproductive traits such as the age of sexual maturity, testes size, sperm characteristics, sexual signals such as antlers, ornamentation, and body size, and behavioral traits such as territoriality, dominance and competition (e.g., Fishman et al., 2022; Mills et al., 2009; Negro et al., 2010; Preston et al., 2012; Thompson et al., 2012). On the other hand, in females, high testosterone levels have been linked to adverse reproductive outcomes, such as increased risk of infertility and cycle disruption, fetal loss, preterm birth, low birth weight, and a high incidence of ovarian dysfunction (Balen et al., 1995; Fuller et al., 1970; James, 2015; Smith et al., 1979; Steinberger et al., 1981).

A heritable component to individual testosterone levels have been shown in both males and females (e.g., Coviello et al., 2011; Flynn et al., 2021; Grotzinger et al., 2018a; King et al., 2004; Kuijper et al., 2007; Mills et al., 2009; Pavitt et al., 2014; Ruth et al., 2020). However, its heritability in free-roaming animals, which are subjected to naturally high environmental variation is understudied (Pavitt et al., 2014).

Given the differential effects of testosterone levels on fitness in males and females, one would expect selection to favor higher testosterone levels in males, and lower levels in females (Moller et al., 2005). A phenomenon, in which alleles affect trait size in both sexes in an opposite manner, so that they promote fitness in one sex and simultaneously decrease fitness in the opposite sex, is referred to as sexual conflict (Bonduriansky and Chenoweth, 2009). This conflict might be resolved by a different genetic architecture between the sexes. Quantitative genetics can be used to assess genetic divergence and the potential for sex-dependent trait evolution. Sex-dependent evolution in a trait shared by males and females is possible through low additive genetic covariance between the sexes, sex differences in additive genetic variances, or both (Cheverud et al., 1985; Lande, 1980; Lynch and Walsh, 1998). The cross-sex genetic correlation between shared male and female traits (rMF) measures the extent of similarity between additive alleles when expressed in both sexes (Bonduriansky and Chenoweth, 2009; Lande, 1980). When rMF is close to one, a shared trait is assumed to be controlled by a common genetic architecture in both sexes. This might constrain one or both sexes from reaching their optimum (Pavitt et al., 2014) despite sex-specific selection. In contrast, when rMF approaches zero, the two sexes differ in genetic architecture for the shared trait or differ in allele expression, allowing the sexes to evolve independently towards trait optimality (Lande, 1980). To date, relatively few studies employed a quantitative genetic approach to understand whether and how the genetic architecture for testosterone levels differs between the sexes, with conflicting findings, that might be attributed to species, age or method differences (Flynn et al., 2021; Grotzinger et al., 2018b; Hoekstra et al., 2006; Pavitt et al., 2014; Ruth et al., 2020; Sinnott-Armstrong et al., 2021). Moreover, while there are a few studies on testosterone heritability in neonates (namely red deer and human) (Caramaschi et al., 2012; Pavitt et al., 2014; Sakai et al., 1991), to the best of our knowledge, there are no studies examining heritability of in utero accumulating testosterone levels, despite the appreciated influence of in utero testosterone exposure on both fetal development and fitness in adulthood (e.g., reviews by Ryan and Vandenbergh, 2002; Zambrano et al., 2014).

Here, we used a quantitative genetic approach to examine the genetic component of in utero accumulated testosterone levels and test whether it differs between the two sexes. Specifically, we assessed i) heritability estimates between parent and offspring within the same sex (i.e., mother-daughter and father-son); ii) heritability estimates between parent and offspring of the opposite sex (i.e., mother-son and father-daughter); and iii) cross-sex genetic correlation. We applied this setup in the nutria (*Myocastor coypus*), a large semi-aquatic rodent native to South America which has a high reproductive potential, including post-gestational estrus, large litters, and year-round receptivity, making it a successful invasive species facing extensive eradication and control efforts worldwide (Carter and Leonard, 2002). It also has a high prevalence of multiple paternity (Fishman et al., 2022). We collected pregnant nutria carcasses obtained during eradication efforts at Agamon Hula Park in Israel and quantified their hair testosterone levels. Unlike saliva and blood concentrations, which are confounded by diurnal variation or hormonal reactivity, hair testing presents long-term steroid accumulation (Grotzinger et al., 2018a; Koren et al., 2002). Furthermore, while most of circulating testosterone is bound to sex hormone binding globulins (SHBG), and thus may not be bioavailable (Dunn et al., 1981), hair testosterone levels reflect free, unbound, and bioavailable testosterone (Chan et al., 2014; Russell et al., 2012; Slezak et al., 2017; Stalder and Kirschbaum, 2012). In the nutria, hair follicles appear at 85–90 days of gestation (around the beginning of the last trimester of their ~132 day-long pregnancy) (Felipe and Masson, 2008; Sone et al., 2008), and have a sufficient amount of hair for testosterone quantification around the last month of gestation, representing accumulating levels of testosterone in the last trimester of pregnancy (Fishman et al., 2019). During that period, nutria fetuses’ hair testosterone levels are not associated with gestational age, nor with maternal testosterone levels (Fishman et al., 2019).

Male nutria fetuses have significantly higher testosterone levels than female fetuses (Fishman et al., 2019). Male fetuses related to fathers monopolizing most or all of the litter paternity have higher testosterone levels and those related to the rare father, while female testosterone levels does not show such relationship (Fishman et al., 2022). These findings are in line with studies that observed higher reproductive success for differentially larger and more aggressive male nutria (Guichón et al., 2003a, 2003b; Túnez et al., 2009). In adults, male body size is strongly associated with testosterone levels, however, this association does not exist in females (Fishman et al., 2022). Moreover, higher testosterone levels in females are related to lower fitness (Fishman et al., 2018). Based on these past findings, we predict that there might be sex-differences in the genetic architecture for testosterone levels in the nutria.

## Methods

### Sample collection

All animals were collected in Agamon Hula Park, in Northern Israel, as a secondary use of culling efforts by authorities in the park during 2013 - 2019. Out of the 142 nutria females dissected, only 16 females had fetuses with sufficient amount of hair for testosterone quantification (gestational age of 110 days or more). Thus, the final sample comprised of 91 fetuses (44 males and 47 females) with gestational ages of 110 - 138 days, belonging to 16 litters. Estimation of gestational age followed Newson’s formula (Newson, 1966) cross-validated with multiple fetal morphometric measurements (Fishman et al., 2018). Fetal sex determination was performed using anatomical and molecular tools as described in (Fishman et al., 2018). Testosterone was extracted and quantified from fetal hair as previously described (Fishman et al., 2019).

### Molecular methods

The molecular analysis was performed on the fetuses and their mothers. Tissue clippings were excised from 91 fetuses and their 16 mothers, preserved in 70% ethanol and refrigerated at −80°C until processing. Total genomic DNA was isolated from 20 mg tissue, and extracted using a Geneaid gSYNC DNA Extraction kit GS100 (Geneaid Biotech Ltd., Taiwan). DNA concentration and cleanness were checked by NanoDrop (Thermo Nanodrop 2000) and diluted as necessary. PCR reactions were performed on the 7 most variable of the 27 microsatellites (Table SI1) developed by (Callahan et al., 2005). PCR reactions contained dye-labeled primers and performed as detailed in (Fishman et al., 2022).

### Genotyping

Fragment analysis was performed on an ABI 3100 Genetic Analyzer. Allelic designations were determined using Peak Scanner Software v1.0. Alleles that were not clear enough for designation were coded as missing data. Validation of the genotyping was done by sporadically rerunning the PCR and the genotyping procedures for various samples. As fetuses were extracted from their mother’s carcass, the assignment of offspring to their mothers was known with 100% certainty. This ‘ground truth’ was used to conduct maternity analysis and resolve mismatches. Allele frequency analysis and maternity analysis were performed using Cervus Software v.3. Full and half-sib analysis and statistical paternity assignment were performed using Colony Software v.01, with 4 repeats of >6 million iterations, followed by a manual check of a sample of litters.

### Statistical analyses

We calculated estimates of heritability for testosterone (log transformed) using the animal model approach following Wilson et al. (2010). We performed Markov Chain Monte Carlo Linear Mixed Models (MCMCglmm) analyses in R ((version 4.0.5, R Core Team 2013, (Hadfield, 2010)). Animal models use pedigree information to partition observed phenotypic variance into different genetic and environmental sources and account for potential confounding effects (fixed factors). We used the animal model to partition the observed phenotypic variance in testosterone (*V*_*P*_), into additive genetic variance (*V*_*A*_) variance due to maternal effects (*V*_*M*_) and residual variance (*V*_*R*_). We used the *us* command to enable estimation of sex-specific additive genetic variance in a trait (females’ additive genetic variance = 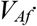 males’ additive genetic variance = *V*_*Am*_), sex-specific maternal effect variance (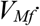,*V*_*Mm*_) as well as an assessment of additive genetic covariance in a trait between males and females (*COV*_*Am*f_) and thus cross-sex genetic correlations (*r*_*mf*_). We constrained the *V*_*R*_ cross-sex covariances to zero using the *idh* command. To define the genetic relatedness between individuals, we constructed a pedigree containing each individual included in the study. Parents of the P (parental) generation were marked as NA because their pedigree was unknown. However, sibling phenotypes also shared effects of the same mother (maternal effects). We estimated narrow-sense heritability in each trait as 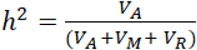 with 95% credible intervals (CIs) for females and males, separately; and calculated whether these differed significantly between the two sexes.

We calculated cross-sex genetic correlation as 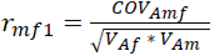 (Lande, 1980). In addition, we were interested in the heritability estimates for all possible parent-offspring combinations: father-daughter 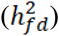, mother-son 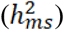, mother-daughter 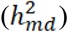, and father-son 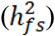. We calculated cross-sex genetic correlation 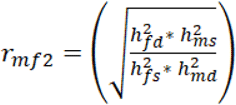 (e.g., Bonduriansky and Rowe, 2005).

All models were run for 100000 iterations, with a burn-in of 2500 iterations, saving samples every 250 iterations. We checked convergence and mixing properties by visual inspection of the chains and checked the autocorrelation values. We ran Heidelberger and Welch’s convergence diagnostics to verify that the number of iterations was adequate for chains to achieve convergence.

## Results

Male fetuses had significantly higher hair testosterone levels than female fetuses (*p* < 0.05). However, male and female fetuses did not differ significantly in their quantitative genetic estimates. The additive genetic variance, *V*_*A*_, accounted for approximately 55% of the observed variance in testosterone levels in both male and female fetuses (%*V*_*A*_, or heritability, is calculated as *V*_*A*_ / *V*_*A*_*+V*_*R*_+*V*_*M*_). The amount of variance due to maternal effects was slightly higher for females than for males (35% and 27%, respectively), and the residual variance accounted for 19% of the variance in males and 12% in females. The amount of additive genetic covariance for testosterone between males and females tended towards zero, while the mean genetic correlation between sexes was estimated to 0.547 (Table 1).

**Table 1.** Estimates of additive genetic variance, variance in maternal effects and residual variance in males and females separately; and additive genetic covariance and genetic correlation between males and females for hair testosterone levels. Males (n=44), females (n=47).

|  | Mean ± 95 % CI | Mean ± 95 % CI |
| --- | --- | --- |
|  | Males | Females |
| **Additive genetic variance** | 0.099 [$<0.001$, 0.177] | 0.108 [0.040, 0.179] |
| **Variance in maternal effects** | 0.048 [$<0.001$, 0.144] | 0.070 [$<0.001$, 0.177] |
| **Residual variance** | 0.034 [0.002, 0.089] | 0.024 [0.002, 0.061] |
| **Heritability** | 0.561 [0.082; 0.962] | 0.557 [0.228, 0.919] |
| **Additive genetic covariance** | 0.055 [-0.011, 0.124] |  |
| **Cross-sex genetic correlation** | 0.547 [-0.034, 0.997] |  |

Similar results were obtained when we calculated heritability between the parents and offspring of the same sex (mother-daughter and father-son) and heritability estimates between the parents and offspring of the opposite sex (mother-son and father-daughter; Figure 1). Heritability from mothers to daughters was estimated to 48% and father to sons’ heritability was estimated to 59% The estimated heritability from mothers to sons was 0.2%, and from fathers to daughters was 0.3%. The mean cross-sex genetic correlation (*r*_*mf2*_) was 0.519. Detailed results are provided in SI2.

**Figure 1.**
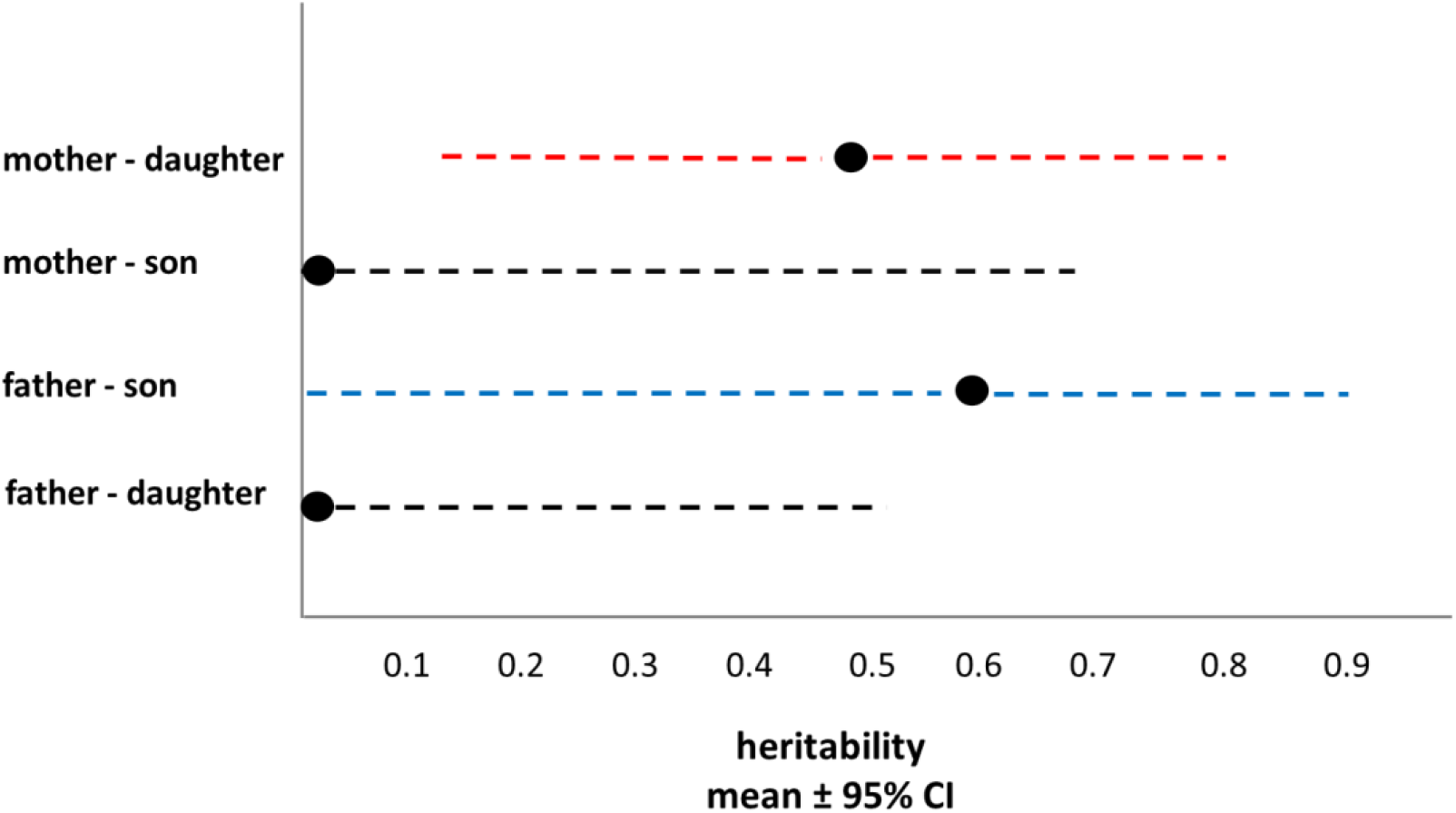
Heritability estimates (mean ± 95% CI) between the parent and offspring of the same sex (mother-daughter and father-son) and heritability estimates between the parent and offspring of the opposite sex (mother-son and father-daughter). Mothers - daughters: 0.477 with 95% CI [0.132, 0.799]; mothers - sons was 0.002 with 95% CI [<0.001, 0.686], father - sons 0.589 with 95% CI [<0.001, 0.892]); fathers - daughters was 0.003 with 95% CI [<0.001, 0.539].

## Discussion

We present the first account of fetal testosterone heritability in a wild population, demonstrating relativity high estimates of heritability between parent and offspring of the same sex and very low heritability estimates between parent and offspring of the opposite sex as well as relatively low cross-sex genetic correlation, implying a potential for sex-specific genetic architectures. We were interested in whether male and female fetuses differ in genetic architecture for testosterone, and thus used an animal model approach to decompose the variance in testosterone levels into additive genetic variance, variance in maternal effects, and residual variance in males and females separately. At the phenotypic level, as expected, male fetuses had higher testosterone levels than female fetuses. Maternal effects explained approximately 30% of both sexes’ variance in testosterone levels. In the uterus, the environmental factors are experienced by the fetus via mediation of the mother, and factors such as the maternal social environment, stress, nutrition and age have been shown to affect offspring testosterone levels (e.g., Da Silva et al., 2001; Kallak et al., 2017; Meise et al., 2016; Ward and Weisz, 1984). The mother may also affect fetal testosterone levels via testosterone transport as observed in avian systems (e.g., Bentz et al., 2016; Schwabl, 1993) or in metabolism of maternal androgens to fetal testosterone as found in hyenas (*Crocuta crocuta*). In rodents, however, this transport may be blocked or metabolized by the placenta, as observed *in vivo* in guinea pigs and rats, but may influence the fetus by affecting the normal placental development and function (Despres et al., 1984; Sathishkumar et al., 2011; Slob et al., 1983; Vreeburg et al., 1981). In the nutria, previous study has shown no association between maternal and fetal hair testosterone levels (Fishman et al., 2019). Maternal effects may also include the intrauterine position of the fetus in relation to same or opposite sex neighbors, which previously shown a relationship with fetal testosterone levels (Fishman et al., 2019). Additive genetic variance explained approximately 55% of the total variance in testosterone levels in both female and male fetuses. Therefore, estimates of heritability (percentage of phenotypic variance explained by additive genetic variance) for testosterone levels were approximately 0.5 in both sexes and did not differ between males (father-son) and females (mother-daughter). However, estimates of heritability between parent and offspring of the opposite sex (mother-son and father-daughter) approached zero. Notably, the estimate of mean additive genetic covariance between males and females was half the additive genetic variance and cross-sex genetic correlation was 0.5, suggesting that the genetic architecture for testosterone differs, at least in part, between males and females.

In theory, a high cross-sex genetic correlation (rMF ~ 1) implies that the sexes largely share the genetic base for a given trait, while low rMF indicates a sex-specific genetic architecture (Lande, 1980). The mean cross-sex genetic correlation (rMF) for testosterone in our study was estimated as 0.5, indicating that the sexes differed to some extent in the genetic basis for the trait. While the 95% CI for rMF was very broad, limiting our conclusions, the estimated heritability between parents and offspring of the opposite sex (mother-son and father-daughter), which was close to zero, suggests that the genetic basis for testosterone may indeed differ between males and females in the nutria. Direct estimations of between-sex genetic correlation of testosterone are relatively rare. In wild red deer (*Cervus elaphus*) calves, Pavitt at al. (2014) found a high genetic correlation between the sexes. A high genetic correlation was also found in humans in 12-year-old twins (Hoekstra et al., 2006), however, in young adults, there was minimal correlation between opposite-sex dizygotic twins salivary testosterone concentrations (Grotzinger et al., 2018b). Recent large-scale genome-wide associations studies (GWAS) in human have shown a significant sex difference in genetic architecture of testosterone and very low between-sex correlation (Flynn et al., 2021; Ruth et al., 2020; Sinnott-Armstrong et al., 2021). Ruth et al. (2020) showed that the genetic component to variation in circulating testosterone concentrations is very different between sexes, with null genome-wide correlations between sexes. Furthermore, they have demonstrated that higher testosterone is harmful for metabolic diseases in women but beneficial in men (Ruth et al., 2020). In their study of sex-specific genetic effects across biomarkers, Flynn (2021) found that between-sex genetic correlations were close to 1 for the majority of the traits examined, while for testosterone the estimated genetic correlation was only 0.12, indicating largely nonoverlapping genetic effects between males and females. Sinnott-Armstrong et al. (2021) also found minimal genetic correlation of testosterone levels between the sexes, in contrast to other biomarkers examined (Sinnott-Armstrong et al., 2021). All three studies demonstrate that in humans, the genetic basis of testosterone is almost completely independent between males and females (Flynn et al., 2021; Ruth et al., 2020; Sinnott-Armstrong et al., 2021).

Though often referred to as a single trait, the use of genetic data from large biobanks have led to the identification of more than a hundred loci associated with factors regulating the biosynthesis, metabolism and excretion of testosterone (Flynn et al., 2021; Ruth et al., 2020; Sinnott-Armstrong et al., 2021). GWAS studies demonstrated that the genetics of testosterone levels is complex and highly polygenic in both males and females (Flynn et al., 2021; Ruth et al., 2020; Sinnott-Armstrong et al., 2021). These studies also show that the regulation of testosterone differs significantly between males and females (Ruth et al., 2020; Sinnott-Armstrong et al., 2021). For example, Sinnott-Armstrong et al. (2021) found that in females, many of the top signals for testosterone are in the steroid bio-synthesis pathway, while a smaller number relate to gonadal development. In males, however, the lead hits for testosterone reflect multiple processes. These differences in the genetic architecture of female and male testosterone are so drastic, that from a genetic point they may be viewed as unrelated traits (Sinnott-Armstrong et al., 2021).

The nutria breeds throughout the year (Fishman et al., 2018; Willner et al., 1979) with no seasonal variation in testosterone levels (Fishman et al., 2018), and an opposite trend of association between testosterone levels and reproductive success between the sexes. Thus, it may be highly beneficial for sons to inherit the high testosterone levels phenotype associated with larger size, which is linked to dominance and reproductive success (Fishman et al., 2022; Guichón et al., 2003a; Túnez et al., 2009), and for daughters to inherit the low testosterone levels phenotype associated with higher females’ fitness (Fishman et al., 2018). Differences in the genetic architecture of the trait may be logical in terms of natural selection: as testosterone has opposite effects on male and female fitness, selection may drive the break of the shared genetic architecture, resolving the sexual conflict. This should allow both sexes to evolve towards their optimum testosterone levels. It is surprising, however, that this sex-differential heritability is expressed already in utero. In human, the heritability estimates of testosterone levels, as well as the between-sex correlation, show large differences across age class and development (e.g., Grotzinger et al., 2018a), possibly corresponding to age-specific differences in biosynthetic pathways (Grotzinger et al., 2018a). For example, at birth, male testosterone levels are higher than females’ (Sakai et al., 1991), decrease to the same level as females at the fifth month after birth (Caramaschi et al., 2012), and increase dramatically at puberty, staying higher when compared to women throughout adulthood (Caramaschi et al., 2012) and references therein). In contrast, nutria pups, which are born highly precocious with a full fur, nails and teeth, reach sexual maturity relatively early, at about 4 months, at a body weight well below the average adult weight (Newson, 1966).

In the fetus, testosterone plays a crucial role in the regulation of growth, maintenance and function of reproductive tissues, impacting the development of the brain, uterus and gonads, both in cell number and morphology, and in receptors regulation (Zambrano et al., 2014). Higher fetal testosterone levels in males compared to females have been observed in various species, for example mice, gerbils, hamsters, rats, nutria, pigs and humans (Clark et al., 1991; Fishman et al., 2019; Ford et al., 1980; Sarkar et al., 2007; Vom Saal and Bronson, 1980; Vomachka and Lisk, 1986; Weisz and Ward, 1980). Male newborns tend to have higher mortality rates (United Nations, 2011) due to a higher vulnerability to perinatal conditions such as prematurity, respiratory distress syndrome and infectious diseases (Waldron, 1998). Fetal testosterone may contribute to this vulnerability via its immunosuppressive effects (Muenchhoff and Goulder, 2014; Olmos-Ortiz et al., 2019), and most notably its competing, or suppressing effects on glucocorticoid production, required for the maturation of multiple fetal systems, including lung maturation (e.g., Morrow et al., 1967; Nielsen et al., 1982; Torday, 1990). Thus, expressing genetic predisposition for higher testosterone levels while in uterus might exert a considerable cost. There is ample evidence that features contributing to male reproductive success, such as tests size and seminal vesicles (Clark et al., 1993, 1990; van der Hoeven et al., 1992; vom Saal, 1989), higher impregnation rates (Clark and Galef Jr, 2000) as well as adult testosterone levels (Clark et al., 1992b; Kilcoyne et al., 2014) are programed and shaped during the prenatal period. Behavioral patterns supporting male reproductive success, such as higher aggressiveness, dominance, territorial behavior, dispersal rates and attractiveness to females are all related to fetal testosterone levels (Clark et al., 1992a, 1990; Drickamer, 1996; Drickamer et al., 1995; Rohde Parfet et al., 1990; vom Saal, 1989; Vom Saal and Bronson, 1978; Zielinski et al., 1992). However, in female fetuses, higher testosterone exposure results in a wide range of fitness reducing effects such as masculinization of internal and external genitalia, delayed puberty onset, altered estrous cycles, increased anovulation, lower number of oocytes, premature loss of fertility, defeminization of GnRHergic synapses, lower receptivity to males and lower attractiveness, (Zambrano et al., 2014 and references therein, Phoenix et al., 1959; Vom Saal and Bronson, 1980). Thus, taken together, it seems highly adaptive that a differential genetic architecture between the sexes would evolve. In light of the early sexual development of the nutria, and its lack of seasonal breeding, which does not promote functional fluctuations in testosterone levels, an early sex-differential expression of the genetic component of testosterone may support both male and female fitness.

