## Supplemental information for "Quantitative genetic study suggests sex-specific genetic architecture for fetal testosterone in a wild mammal"

Table SI1. Microsatellite primers used for assessing multiple paternity in 91 *Myocastor coypus* fetuses. The number of alleles ($N_A$), observed ($H_O$) and expected heterozygosity ($H_E$), and polymorphic information content (PIC). Loci that are not in Hardy-Weinberg equilibrium (HWE) are marked with an asterisk.

| Locus | Primer sequence 5'-3' | N | $N_A$ | $H_O$ | $H_E$ | PIC |
|---|---|---|---|---|---|---|
| McoD214 | F: TTCACAAATCAGAGGCTACAATC<br>R: GTGTTCTTCATTTGATGCTCAG | 385 | 10 | 0.865 | 0.823 | 0.798 |
| McoA02 | F: CCCACATGTATTTGCTTTTGAG<br>R: CAGATTCTGAGCACAGTGAGAC | 385 | 5 | 0.657 | 0.648 | 0.613 |
| McoD217 | F: AGTCCAATCTACATAGCAAGGC<br>R: AGATAGCCCAGCAGAATAAGTG | 366 | 3 | 0.454 | 0.448 | 0.398 |
| McoD10* | F: TTTTGATACTAGCACCAATTATCTTTT<br>R: CTAAAGTTTGCAGCCTTGATTC | 373 | 6 | 0.622 | 0.688 | 0.633 |
| McoD60* | F: TCACTCAATAAATACTCAGGATGC<br>R: TGGGTGTAGGTGTAGGTAGACA | 379 | 5 | 0.752 | 0.726 | 0.680 |
| McoD215 | F: GTTCAGACTTAGGAGTTGCTGG<br>R: GCTCCCTCGATACATTGATTAG | 381 | 7 | 0.446 | 0.449 | 0.427 |
| McoD69* | F: TTCCATCCCTGGTACCATATAC<br>R: TGAAGCATTAGATGCCTTTGTA | 366 | 3 | 0.516 | 0.606 | 0.537 |
| | | Average | 5.57 | 0.635 | 0.627 | 0.584 |

**SI2. Detailed results for "Quantitative genetic study suggests sex-specific genetic architecture for fetal testosterone in a wild mammal"**

Male fetuses had significantly higher hair testosterone levels than female fetuses (post. mean difference (males *minus* females) = 0.217; 95% CI [0.017, 0.414]; p = 0.032). However, females and males did not differ significantly in their quantitative genetic estimates; the amount of additive genetic variance (females: post. mean = 0.108; 95% CI [0.040, 0.179]; males: post. mean = 0.099; 95% CI [<0.001, 0.177]; post. mean difference = 0.009; 95% CI [-0.101, 0.120], the amount of variance due to maternal effects (females: post. mean = 0.070; 95% CI [<0.001, 0.177]; males: post. mean = 0.048; 95% CI [<0.001, 0.144]; post. mean difference = 0.022; 95% CI [-0.119, 0.051]), and the amount of residual variance (females: post. mean = 0.024; 95% CI [0.002, 0.061]; males: post. mean = 0.034; 95% CI [0.002, 0.089]; post. mean difference = -0.010; 95% CI [-0.081, 0.052]). The heritability of testosterone was estimated to posterior mean of 0.557 with 95% CI [0.228, 0.919] in females, and to posterior mean of 0.561 with 95% CI [0.082; 0.962] in males. Heritability estimates did not significantly differ between the two sexes (=-0.004; 95% CI [-0.531, 0.556]). Nevertheless, the amount of additive genetic covariance for testosterone between males and females tended towards zero (post. mean =0.055; 95% CI [-0.011, 0.124]). The mean genetic correlation between sexes was estimated to 0.547 with 95% CI [-0.034, 0.997].

Similar results were obtained when we calculated heritability between the parents and offspring of the same sex (mother-daughter and father-son) and heritability estimates between the parents and offspring of the opposite sex (mother-son and father-daughter; Figure 1). Heritability from mothers to daughters was estimated to the posterior mean 0.477 with 95% CI [0.132, 0.799]; whereas father to sons' posterior mean heritability was 0.589 (95% CI [<0.001, 0.892]). The estimated posterior mean heritability from mothers to sons was 0.002, 95% CI [<0.001, 0.686], and from fathers to daughters was 0.003, 95% CI [<0.001, 0.539]. The mean cross-sex genetic correlation ($r_{mf2}$) was 0.519 with 95% CI [<0.001, 0.864].
